# Genetic basis and selection of glyceollin induction in wild soybean

**DOI:** 10.1101/2022.12.17.520864

**Authors:** Farida Yasmin, Hengyou Zhang, Larry Leamy, Baosheng Wang, Jason Winnike, Robert W. Reid, Cory R. Brouwer, Bao-Hua Song

**Affiliations:** Department of Biological Sciences, The University of North Carolina at Charlotte, NC 28223, USA; Key Laboratory of Plant Resources Conservation and Sustainable Utilization, South China Botanical Garden, Chinese Academy of Sciences, Guangzhou 510650, China; Center of Conservation Biology, Core Botanical Gardens, Chinese Academy of Sciences, Guangzhou 510650, China; David H. Murdock Research Institute, NC Research Campus, Kannapolis, NC 28081, US; Department of Bioinformatics and Genomics, The University of North Carolina at Charlotte, NC 28223, USA

**Keywords:** Epistasis, Gene cluster, mGWAS, phytoalexin, Plant and human health, Selection, Transcription factors, Wild soybean

## Abstract

- Glyceollins, a family of phytoalexin induced in legume species, play essential roles in responding to environmental stresses and in human health. However, little is known about the genetic basis and selection of glyceollin induction.
- We employed a metabolite-based genome-wide association (mGWA) approach to identify candidate genes involved in glyceollin induction from genetically diverse and understudied wild soybeans subjected to soybean cyst nematode stress.
- Eight SNPs on chromosomes 3, 9, 13, 15, and 20 showed significant association with glyceollin induction. Six genes close to one of the significant SNPs (ss715603454) on chromosome 9 fell into two clusters, and they encode enzymes in the glycosyltransferase class within the phenylpropanoid pathway. Transcription factors (TFs) genes, such as *MYB* and *WRKY* were also found within the linkage disequilibrium of the significant SNPs on chromosome 9. Epistasis and a strong selection signal were detected on the four significant SNPs on chromosome 9.
- Gene clusters and transcription factors may play important roles in regulating glyceollin induction in wild soybeans. Additionally, as major evolutionary factors, epistatic interactions and selection may influence glyceollin variation in natural populations.

## Introduction

Plants produce diverse specialized metabolites (also known as secondary metabolites or phytochemicals), which play a vital role in adapting to changing environments. Phytoalexins are specialized metabolites synthesized *de novo* in response to various biotic and abiotic stresses. Examples include indole alkaloid camalexin in *Arabidopsis,* phenolic aldehyde gossypol in cotton, phenylpropanoid stilbenes in grapevines, isoflavonoid-derived glyceollins in legume, and momilactones and phytocassanes terpenoids in rice (Donnez et al., 2011, Jahan et al., 2019, Jeandet et al., 2002, Jeandet et al., 2020, Saga et al., 2012, Wang et al., 2009, Yamamura et al., 2015). Isoflavonoids have become a research hot spot due to their various pharmacological properties and essential roles in plant defense. The major isoflavones in soybeans are genistein, daidzein, and glycitein, and they make up about 50%, 40%, and 10%, respectively, of the total isoflavone content. Trace amounts of glyceollins are induced transiently with abiotic and biotic stresses (Jahan et al., 2019, Subramanian et al., 2006). They have multiple effects, including fostering symbiosis between soybean and *Bradyrhizobium japonicum* and inhibiting the growth of various microbes (Graham and Graham, 1996, Subramanian et al., 2006). Moreover, they have anti-cancer, antioxidant, and neuroprotective properties (Bamji and Corbitt, 2017, Kim et al., 2012, Nwachukwu et al., 2013, Seo et al., 2018). However, studies on glyceollins are mainly focused on their medicinal properties, while little is known about how their induction is regulated.

Phytoalexins have been considered the target of natural selection due to their activities in biotic and abiotic stress responses in natural environments (Miyamoto et al., 2016, Pichersky and Gang, 2000, Qi et al., 2004). Therefore, in our study, we chose wild soybean (*Glycine soja*), a wild relative of soybean (*Glycine max*), to delineate genetic basis and evolution of glyceollin accumulation resulting from biotic stress, i.e., soybean cyst nematode (SCN), the most devastating soybean pest worldwide (Tylka and Marett, 2021). Wild soybeans thrive in diverse habitats and harbor much higher, underexplored genetic diversity than cultivated soybean (Zhang et al., 2019). Hence, it is an ideal system to understand the genetic basis and evolution of glyceollin variation. Eventually, the essential genes identified in wild soybean can be used for metabolic engineering or in a breeding program to develop nutrition-rich biofortified soybean cultivars as they exhibit similar genome size and content with small reproductive isolation (Singh and Hymowitz, 1999).

A metabolic gene cluster is a group of (two or more) genomically co-localized and potentially coregulated non-homologous genes that encode enzymes involved in a particular metabolic pathway (Nützmann et al., 2016, Töpfer et al., 2017). They have been a common phenomenon since the early days of microbial genetics (Koonin, 2009, Rocha, 2008, Zheng et al., 2002). However, gene clusters in plant metabolic pathways have been discovered only recently, even though microbes and plants are both extremely rich sources of metabolic diversity. A study by Chae et al. (2014) on metabolic gene clusters in *Arabidopsis*, soybean, sorghum, and rice suggested that approximately one-third of all the metabolic genes in *Arabidopsis*, soybean, and sorghum, and one-fifth in rice were rich in gene clusters across primary and specialized metabolic pathways (Chae et al., 2014). There is compelling evidence indicating that the highly plastic plant genome itself generates metabolic gene clusters via gene duplication, neofunctionalization, divergence, and genome reorganization instead of horizontal gene transfer from microbes (Osbourn and Field, 2009). This suggests that plants rewire their genome to gain new adaptive functions driven by the need to survive in distinct environments. Systematic mining and functional validation of the candidate genes in such clusters will facilitate the discovery of new enzymes and chemistries that render pathway prediction. Moreover, metabolic gene clusters are likely to be located within dynamic chromosomal regions, and thus, many identified so far may be due to recent evolution (Field et al., 2011, Matsuba et al., 2013, Qi et al., 2004). If so, investigation of these clusters can provide insights into their evolutionary history. The vast and diverse array of specialized metabolites that are produced through multi-step metabolic pathways play an important role in plant adaptation to various ecological niches. However, the occurrence, prevalence, and evolution of such gene clusters in plants are largely unknown. Thus, the study of plant metabolic gene clusters has implications for molecular biology and evolutionary genomics (Chavali and Rhee, 2018, Nützmann et al., 2016, Takos and Rook, 2012, Yeaman and Whitlock, 2011).

Due to the extraordinary metabolic diversity, to date, less than 50 plant-specialized metabolic pathways have been biochemically and genetically identified (Nützmann et al., 2016). Metabolomic GWAS (mGWAS) offers an effective approach to understand the genetic basis of metabolites and their associated traits (Chan et al., 2010, Chan et al., 2011, Luo, 2015, Riedelsheimer et al., 2012). mGWAS allows the identification of common polymorphic regions controlling complex metabolic traits by substantially increasing association panel and genome-wide molecular markers. Besides elucidating genetic architecture, mGWAS can also be used to infer gene functions (Luo, 2015). Hence, mGWAS provides a comprehensive approach to discovering candidate genes. Thus far, it has been used to uncover the genetic basis of variations of a number of different metabolites. For example, Chen et al. (2014) carried out a rice mGWAS study that identified 36 candidate genes influencing the variation of metabolites with physiological and nutritional importance (Chen et al., 2014).

The isoflavonoid pathway has been relatively well studied (Sukumaran et al., 2018). However, it is still not clear how glyceollin induction is regulated. This study is the first to employ genomic and evolutionary approaches to understand the genetic basis and selection of glyceollin induction. Our study provides a fundamental basis for the long-term goal of developing glyceollin-fortified soybean cultivars that would improve plant and human health to meet current and future global challenges. In this study, we aim to address these three questions: (1) What is the genetic basis of variation in glyceollin induction by SCN? (2) Are there any gene clusters and transcription factors involved in glyceollin variation? (3) Are epistatic interactions and natural selection important evolutionary factors influencing the variation of glyceollin induction?

## Materials and Methods

### Plant materials

A total of 264 accessions of wild soybean, *Glycine soja*, from a wide geographic range, originally collected from China, Japan, Russia, and South Korea, were utilized (Table S1). The seeds of these ecotypes were obtained from the USDA national germplasm resources laboratory (https://www.ars-grin.gov/).

### Plant preparation, SCN inoculation, and sample collection

Seed preparation, germination, transplanting, and soybean cyst nematode (SCN, *Heterodera glycines Ichinohe*, HG type 1.2.5.7) inoculation were performed following a previously developed protocol (Zhang et al., 2017a, Zhang et al., 2017b, Zhang and Song, 2017). Whole root tissues were collected and weighed five days post-infection (dpi). The 5 dpi time point was chosen because our previous study suggested a significant inhibition in SCN development in a resistant genotype compared to normal growth in a susceptible genotype (Zhang et al., 2017a, Zhang et al., 2017b). All samples were flash frozen in liquid nitrogen and stored at −80 °C. Four biological replicates per wild soybean genotype were used, eventually a total of 1,020 samples.

### Metabolite extraction and quantification

We employed the extraction method of metabolites from root tissue described in Strauch et al., (2015). The metabolite profiling was provided by the service from David H. Murdock Research Institute at the North Carolina Research Campus. Peaks that were consistently detected in at least three biological replicates within each genotype were used for downstream analyses. Each metabolite was confirmed using pure standard compounds, including daidzein, daidzein-d6, and glyceollin. Due to the low concentrations of these compounds and the small sample masses of the wild soybean root samples that had been collected, we used a signal-to-noise ratio of ≥ 10 for the measurement of the peaks for glyceollin and daidzein. Our method successfully measured daidzein (μg/g root) and glyceollin (unitless) in 264 accessions of wild soybean *G. soja* roots quantitatively and semi-quantitatively, respectively. Following method development, optimization, and analyses of the test samples, calibration curves were designed using at least six different concentrations of daidzein, created in triplicate to quantify known concentrations of daidzein and glyceollin. A second-degree polynomial was derived from the known concentrations of the standard curve samples and the mass spectrometer response (daidzein/internal standard) from the standard curve data. The resulting polynomial was used to calculate the concentrations of daidzein in the experimental samples. Low, medium, and high QC (quality control) samples were created to assess the accuracy of the calculations. We used the ratio of glyceollin (unitless) to daidzein (μg/g root) (GVSD) as our phenotypic trait. This phenotype henceforth is denoted GVSD.

### Genotypic data

Genotype data for the 264 accessions were obtained from SoySNP50K (Song et al., 2013), which included 32,976 genome-wide single nucleotide polymorphic markers (SNPs) with a minor allele frequency (MAF) of at least 5%.

### Metabolite-based genome-wide association study (mGWAS) and linkage disequilibrium estimation

Our genome-wide association analysis was conducted on GVSD (a ratio of glyceollin mean to daidzein mean) in response to SCN infection on all 264 ecotypes using the BLINK algorithm implemented in the GAPIT R package (2.0) (Tang et al., 2016). To minimize false-positive associations, we controlled population structure among genotypes with four principal components. Heritability estimate and SNP effect were calculated by running GWAS applying CMLM and MLM methods, respectively, implemented in the GAPIT R package (2.0) (Tang et al., 2016).

A conventional Manhattan plot was generated using the qqman R package to visualize the SNPs (Turner, 2014). In addition to the genome-wide significant threshold, we also calculated the chromosome-wide Bonferroni thresholds using independent SNPs estimated on each chromosome following the method of Li and Ji (2005) (Li and Ji, 2005). Linkage disequilibrium (LD) was calculated across the panel with the TASSEL program, version 5 [6], for the significant SNPs identified from the GWAS analysis. LD was measured using squared correlation R-squared (r^2^) of 0.2 (upper right in the LD plot) and *p*-value < 0.05 (the lower left in the LD plot). A pairwise LD was generated following the R function described by Shin et al. (2006) (Shin et al., 2006). Genes within LD blocks containing significant SNPs were identified as potential sources of candidates for further analyses.

### Identification of candidate genes

For extensive gene mining of our identified gene pool, we used an array of bioinformatics tools. Such an approach can improve the accuracy of candidate gene and gene cluster predictions and resolve inconsistencies among the bioinformatics tools (Chavali and Rhee, 2018). Specifically, a pairwise linkage disequilibrium (LD) analysis was initially used for potential candidate gene identification. Then, genes in each LD block were examined as potential candidate genes, and their annotations were obtained from the Phytozome v13 database (Goodstein et al., 2011). Afterward, a GO enrichment analysis of the identified candidate genes was performed using ShinyGO v0.66: Gene Ontology Enrichment Analysis (*p*-value cutoff (FDR, false discovery rate) = 0.05) (Ge et al., 2020), Soybase GO Enrichment Data (Grant et al., 2010). To investigate the involvement of these potential candidate genes in metabolic pathways, a database search was performed through an annotation file from Phytozome v13 (Goodstein et al., 2011), Soybase (Grant et al., 2010), SoyCyc 10.0 Soybean Metabolic Pathway (Hawkins et al., 2021), and Pathview databases (Luo et al., 2017). Finally, a PMN plant metabolic cluster viewer was applied to categorize enzymes into classes (signature or tailoring) and metabolic domains (Hawkins et al., 2021).

### Analysis of epistatic interactions

For any significant SNPs uncovered in the GWAS analysis, it is useful to test whether, beyond their direct effects, they also exhibited interactive effects on GVSD. To accomplish this, we first produced numerically formatted genotypes, in which the homozygous genotype index value is 1 and −1 and the heterozygous 0. This allows us to test for epistasis for each pairwise combination in a simple general linear model with 1 degree of freedom for the additive effects of each of the two SNPs and their interaction. We included the first four principal components from the GAPIT analysis in the model to be consistent with the GWAS scan, where these components were used to adjust for structural relatedness (see below). The significance of all interactions was evaluated with the sequential Bonferroni procedure. To illustrate the interactions of SNP pairs, we also calculated regressions of GVSD on each SNP, but at each of the three genotypes (using the −1, 0, and 1 index values) of the second SNP involved in the significant interaction.

### Extended haplotype homozygosity analyses

To test allele-specific selection patterns of the identified significant SNPs, we analyzed extended haplotype homozygosity (EHH, (Sabeti et al., 2002)) for each significant SNP. The EHH analysis was conducted in SELSCAN v.1.2.0a (Szpiech and Hernandez, 2014) with default parameters, and only SNPs with MAF > 0.05 was used in this analysis.

## Results

### Genomic dissection of glyceollin accumulation upon biotic induction

We identified a total of eight significant SNPs, with four located on chromosome 9 and the others on chromosomes 3, 13, 15, and 20 (Fig. **1a**, Table 1). These SNPs were identified based on both genome-wide Bonferroni threshold of 5.104 and chromosome-wide Bonferroni thresholds that varied narrowly from 3.79 to 3.82 among the 20 chromosomes (3.803 on chromosome 9) (Figs **1a,b**, Table S2). The manhattan and Q-Q (quantile-quantile) plots are shown in Fig.**1a,b,c**. The four significant SNPs on chromosome 9 are located close to each other within a 535 kb region (Table S2). The broad-sense heritability (*h^2^*) was estimated 35% (Table S2).

**Fig. 1.**
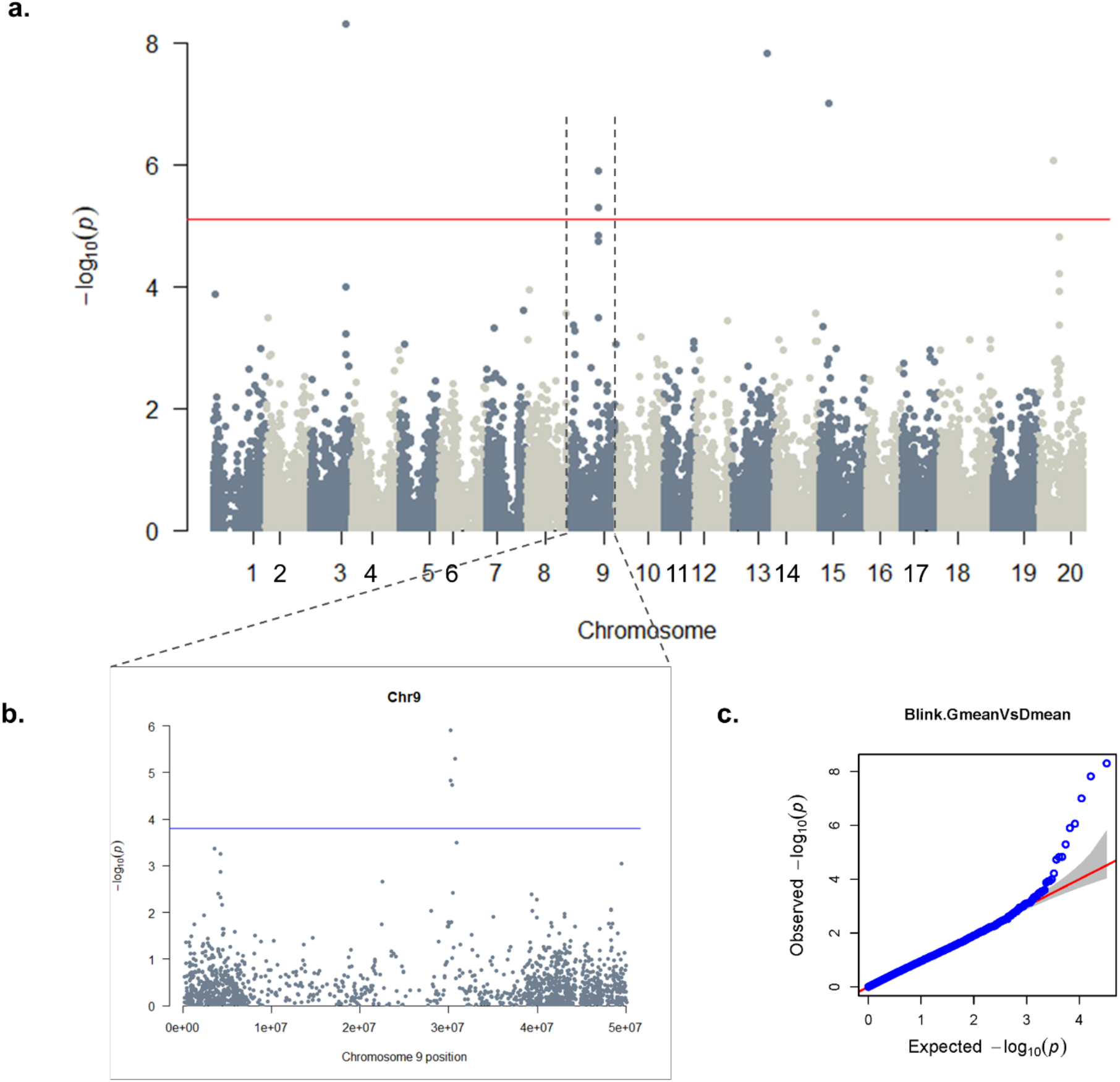
GWAS of Glyceollin induction with SCN stress: A genome-wide **(a)**and chromosome-wide **(b)**Manhattan plots, with thresholds of 5.104 and 3.803, respectively; **(c)**quantile-quantile (QQ) plot. Significant SNPs are found on chromosomes 3, 9, 13, 15 and 20 at a 5% genome-wide threshold, the probability of 7.86×10^-6^ resulted in a threshold of 5.01 (solid red line in the genome-wide Manhattan plot) **(a)**. The 5% chromosome-wide LOD threshold resulted in significant p-values of 1.57×10^-4^ (threshold 3.803, solid blue line) **(b)**.

**Table 1.** Identification of significant SNPs and functional annotation of the plausible candidate genes.

| Significant SNP | Chromosome | Functional annotation of associated genes |
| --- | --- | --- |
| ss715585948 | Gm03 | *WRKY* family transcription factor family protein<br>Zinc fingers superfamily protein |
| ss715603454 | Gm09 | UDP-glucosyl transferase 88A1 |
| ss715603455 | Gm09 | *RING/U-box* superfamily protein, *RING/FYVE/PHD* zinc finger superfamily protein |
| ss715603462 | Gm09 | *WRKY* family transcription factor family protein<br>*MYB* domain |
| ss715603471 | Gm09 | Zinc fingers superfamily protein<br>*Cytochrome P450* enzyme family<br>Zinc finger, *RING-type*; Transcription factor<br>jumonji/aspartyl beta-hydroxylase |
| ss715615975 | Gm13 | *bZIP* transcription factor<br>*RING/U-box* superfamily protein, *RING/FYVE/PHD* zinc finger superfamily protein<br>Zinc fingers superfamily protein<br>*NAC* transcription factors<br>*Cytochrome P450* enzyme family |
| ss715620269 | Gm15 | *RING/U-box* superfamily protein, *RING/FYVE/PHD* zinc finger superfamily protein<br>*WRKY* family transcription factor family protein<br>*MYB* domain |
| ss715636844 | Gm20 | UDP-Glycosyltransferase superfamily protein<br>UDP-glucosyl transferase 85A2<br>hydroxy methylglutaryl CoA reductase 1<br>*Cytochrome P450*, family 71, subfamily B, polypeptide 34<br>cytochrome p450 79a2<br>*RING/U-box* superfamily protein, *RING/FYVE/PHD* zinc<br>finger superfamily protein<br>Zinc fingers superfamily protein |

### Linkage disequilibrium analysis and candidate gene identification

We identified a total of 666 possible candidate genes within the linkage disequilibrium (LD) blocks of the eight significant SNPs (soybean reference genome *Glycine max* Wm82.a2.v1) (Goodstein et al., 2011, Zhou et al., 2015). The LD block on chromosome 9 showed the strongest LD with a long range compared to the others (Figs **2b**, **S1, S2**). We considered r^2^>0.2 as a cutoff for our LD analysis, where r^2^ is the extent of allelic association between a pair of sites (Weir, 1990). Candidate gene *Glyma.09G128200* shows the highest level of LD near the significant SNPs on chromosome 9 compared to the LD block for the rest of the significant SNPs on this chromosome (Figs **2b**, **S1**). The functional annotation of the candidate genes within this block is biosynthetic enzymes involved in isoflavonoid pathway, as well as regulatory genes such as *WRKY* and *MYB* transcription factors (Tables 1, S3, and S4), which may indicate their transcriptional level involvement in glyceollin induction in response to SCN stress (Colinas and Goossens, 2018).

**Fig. 2.**
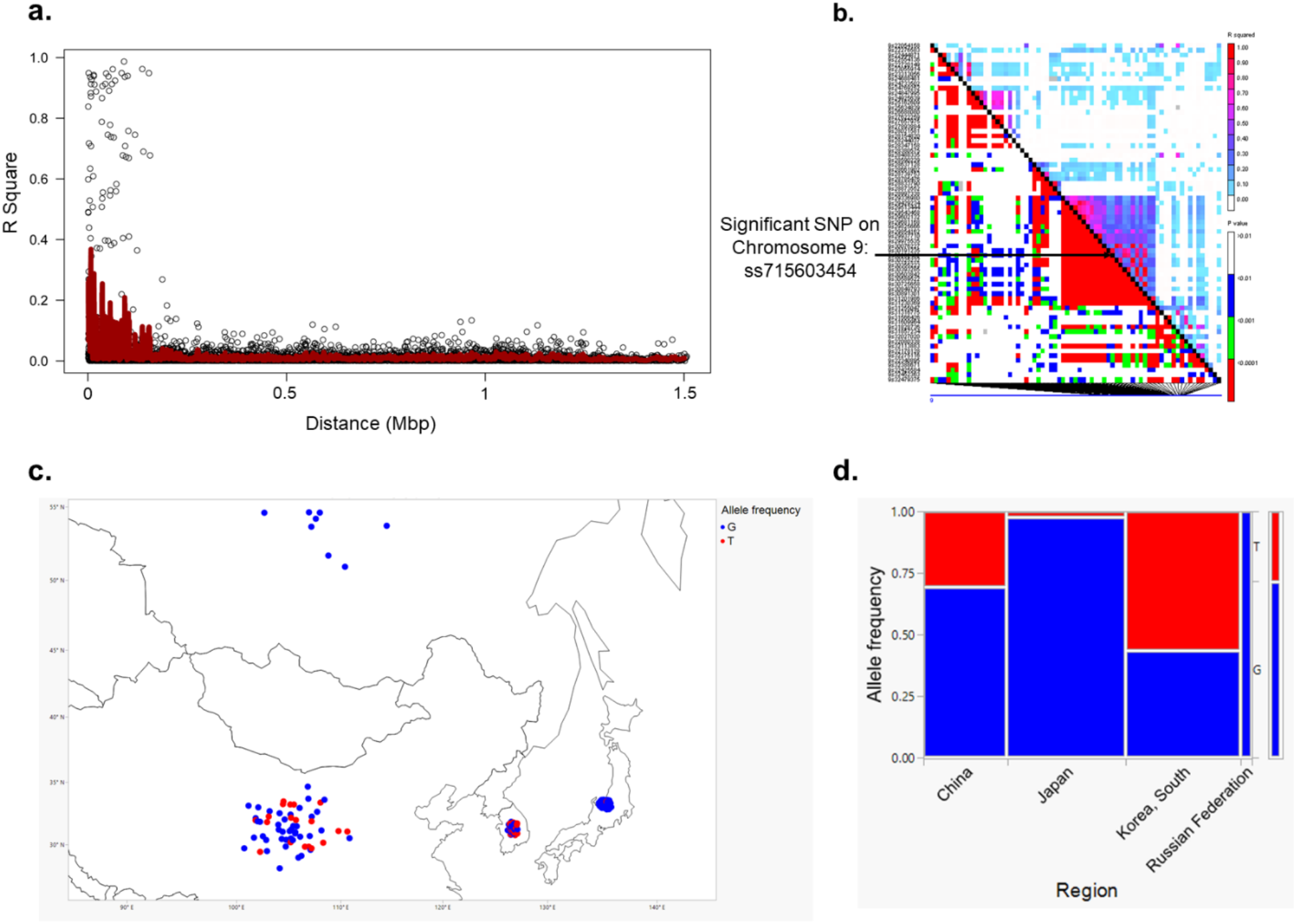
An LD decay measured as R square for pairwise markers and plotted against their distance **(a)**and LD plot for chromosome 9 for significant SNPs. The black diagonal denotes LD between each site and itself **(b)**. Geographic range of the alleles of significant SNPs close to the gene clusters on chromosome 9 **(c)**. Allele frequency in each population. Allele frequency in different geographic regions for a significant SNP was generated using JMP®, Version 15. SAS Institute Inc., Cary, NC, 1989-2021. **(d)**.

We also found putative genes encoding enzymes involved in the specialized metabolic pathways within the LD blocks of the significant SNPs on chromosomes 3, 13, 15, and 20. The enriched GO category includes flavonoid biosynthesis pathway, phenylpropanoid metabolic process, linamarin biosynthesis, and terpenoid biosynthesis (Table S5). Apart from the biosynthetic enzymes on these chromosomes, we also found transcription factor genes, such as *WRKY, MYB*, and *NAC* (Table S5).

### Metabolic gene clusters identification

We were particularly interested in the candidate genes in the branch from daidzein to glyceollin in the isoflavonoid biosynthesis pathway (Lozovaya et al., 2007). We found that the identified candidate genes on chromosome 9 are clustered together, and they fell into two clusters. Both of these two clusters belong to tailoring enzyme glycosyltransferase within phenylpropanoid specialized metabolic domain. And six genes are within the branch of isoflavonoid biosynthesis pathway. Two of these six genes, *Glyma.09G127200* and *Glyma.09G127300*, are called cluster 1, while the rest four (*Glyma.09G127700, Glyma.09G128200, Glyma.09G128300*, and *Glyma.09G128400*) are called cluster 2 (Table S3).

Further investigation of annotation of these candidate genes within the gene clusters (Table S4), we found *Glyma.09G127200* gene encodes a glucosyltransferase that may act on 4’-methoxy isoflavones biochanin A, formononetin, 4’-hydroxy isoflavones genistein, and daidzein substrates. However, the enzyme does not act on isoflavanones, flavones, flavanones, flavanols, or coumarins (Köster and Barz, 1981). Within the same cluster, *Glyma.09G127300* has similar annotations and functions as *Glyma.09G127200.* Interestingly, the four genes within cluster 2 have a similar functional annotation as *Glyma.09G127200 and Glyma.09G127300* in cluster 1, and all these four genes encode isoenzymes (Table S4). Such a link between these two gene clusters indicates their proximity in the metabolic pathway.

### Epistatic interactions among all significant SNPs

The results of the epistasis tests for each of the 28 pairwise combinations of the eight significant SNPs are shown in Table 2. Three probabilities, all associated with the SNP on chromosome 20, were not estimable (Table 2). Among the remaining 25 SNP pairs, 20 show statistical significance. Particularly noticeable is the high significance for all interactions of the SNPs on chromosomes 3, 13, and 15. Three of the six pairs among the four SNPs on chromosome 9, all involving ss715603462, also are statistically significant. In general, therefore, this is evidence for substantial epistasis among these SNPs affecting GVSD.

**Table 2.**
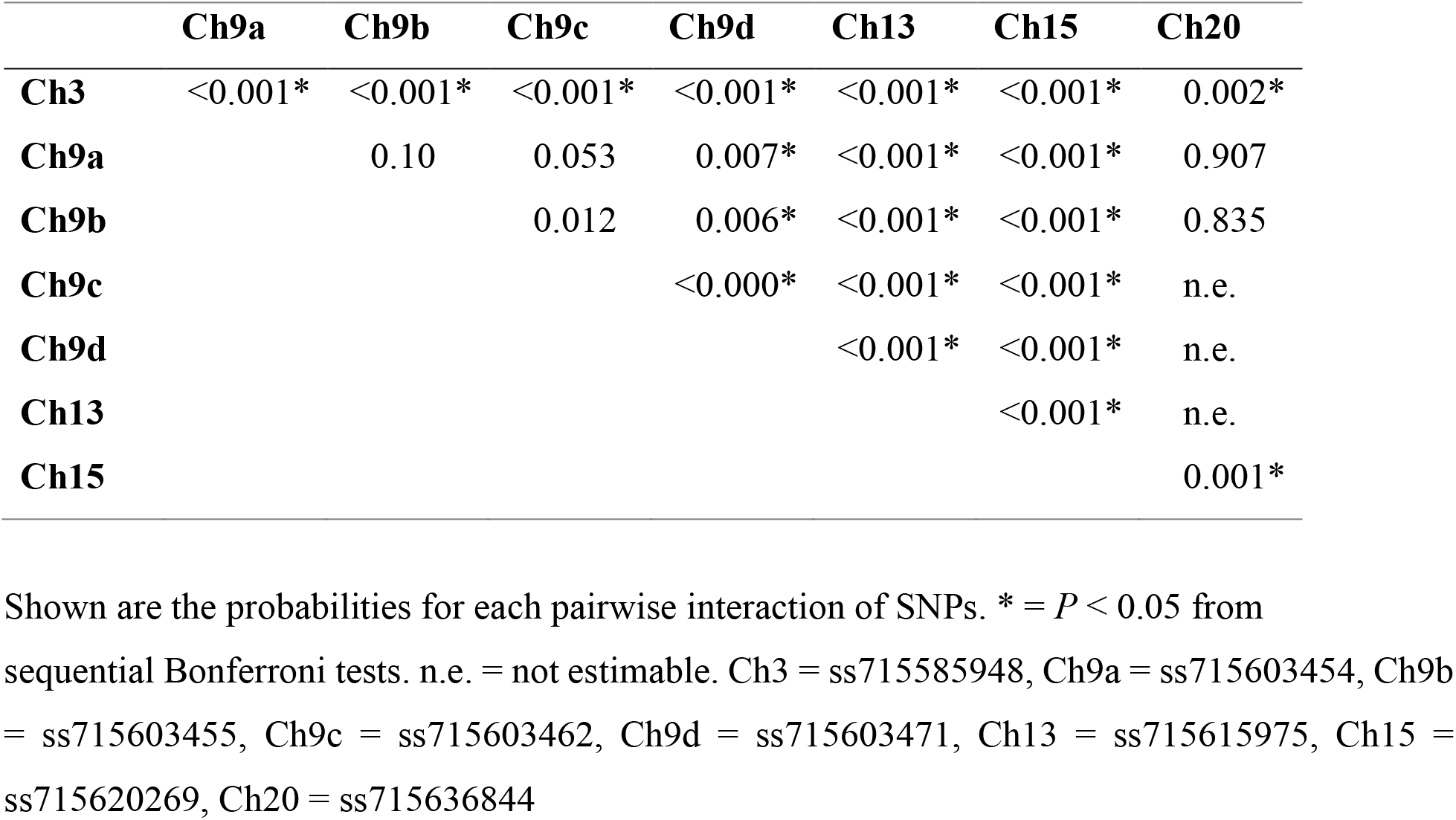
Epistasis for the eight significant SNPs.

These epistatic interactions of the SNP pairs are illustrated in Fig. **3** for each of the four chosen combinations. For example, in panel **a** (Fig. **3a**), it can be seen that regression slopes of GVSD on ss715603454 are close to 0 for ss71585948 CC genotype but are positive for TC and especially TT genotypes. In panel d (Fig. **3d**), regression slopes of GVSD on ss715603471 are negative for ss715603462 AA and GA genotypes but positive for GG genotypes. With no epistasis, these slopes would be expected to be roughly parallel, but in fact, they diverge considerably from parallelism in these four examples.

**Fig. 3.**
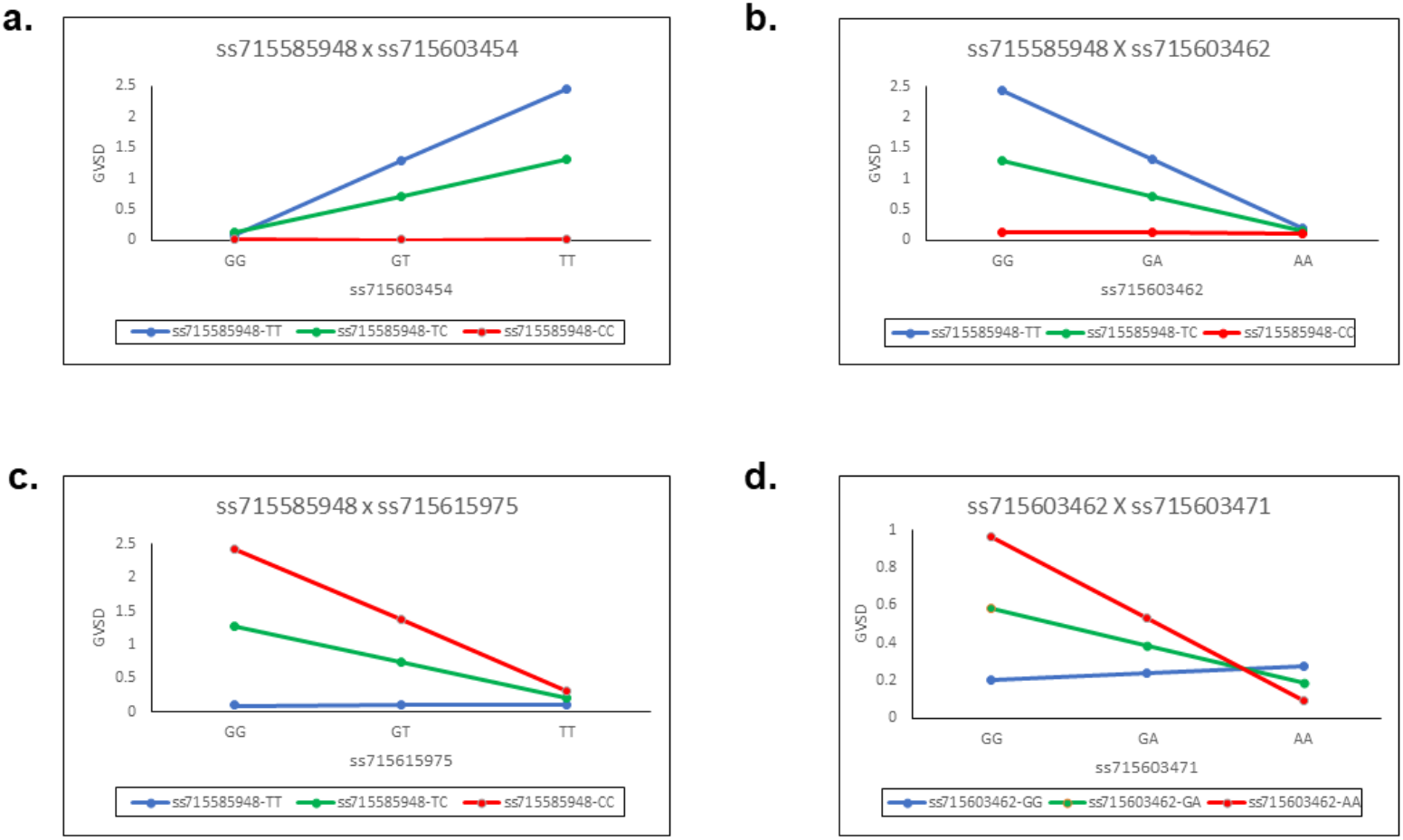
Epistatic interactions of the SNP pairs for each of four chosen combinations. Regression slopes of GVSD on ss715603454 are close to 0 for ss715603454 CC genotypes but are positive for TC and especially TT genotypes **(a)**. Regression slopes of GVSD on ss715603462 are close to 0 for ss715585948 CC genotypes but are negative for TC and especially TT genotypes **(b)**. Regression slopes of GVSD on ss715615975 are close to 0 for ss715585948 TT genotypes but are negative for TC and especially CC genotypes **(c)**. Regression slopes of GVSD on ss715603471 are negative in sign for ss715603462 AA and GA genotypes, but positive in sign for GG genotypes **(d)**.

### Significant SNPs exhibited extended haplotype homozygosity

The extended homozygosity analysis (EHH) analyses revealed allele-specific EHH values of the significant SNPs (ss715603454, ss715603455, ss715603462, and ss715603471) on chromosomes 9 (**Fig. 4**). For example, T allele of ss715603454 showed much higher EHH value than G allele. Alleles of significant SNPs on the other chromosomes showed compatible EHH values (**Fig. 4**).

**Fig. 4.**
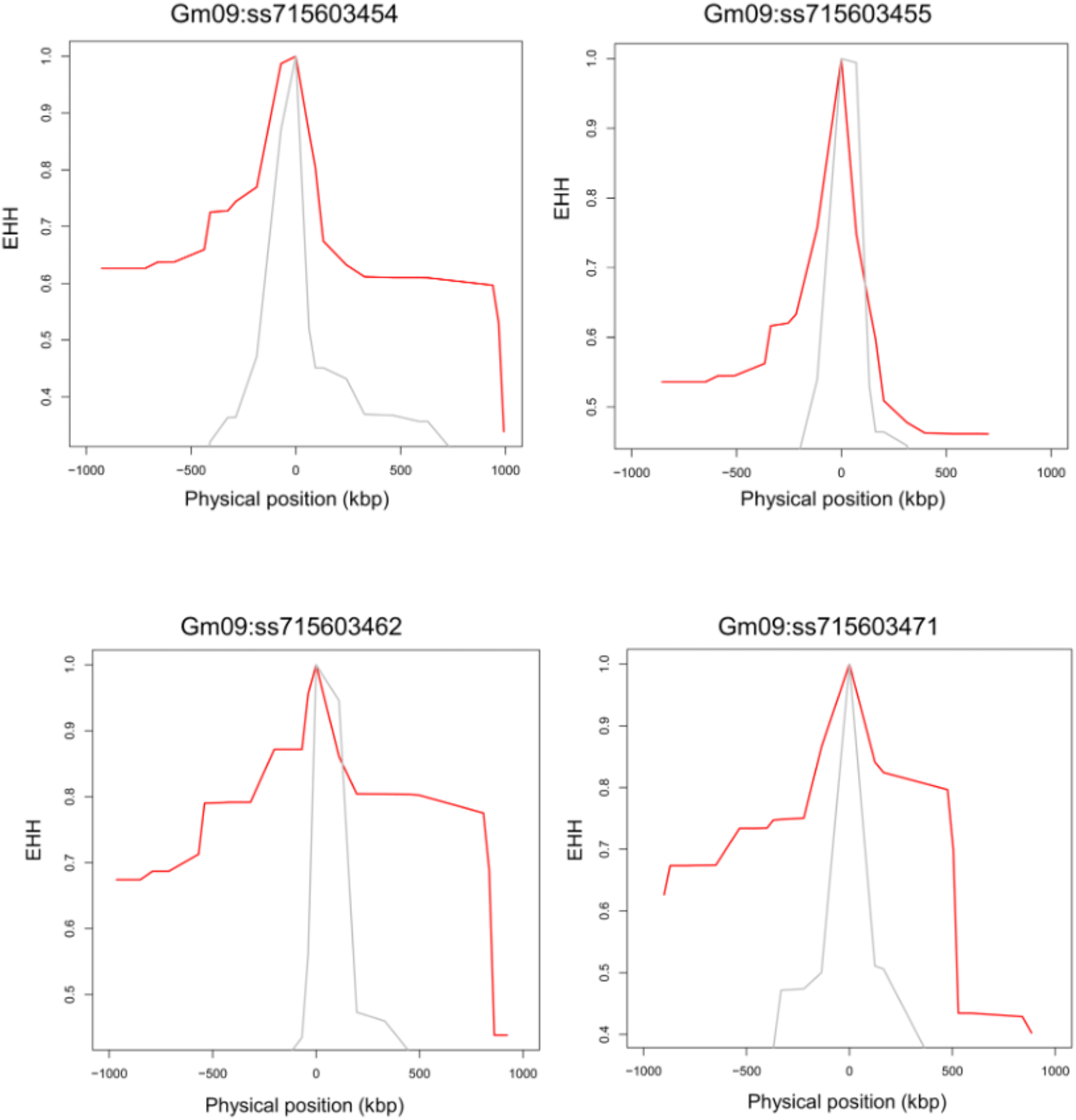
Allele-specific Extended Haplotype Homozygosity (EHH) for four significant SNPs on chromosomes 9.

## Discussion

### Metabolic gene clusters in glyceollin induction

Gene clusters have been reported to play important roles in phytochemical diversity in *Arabidopsis*,sorghum, soybean, and rice (Chae et al., 2014). However, their roles in regulating metabolic variation in wild species are relatively less investigated. Even though the isoflavonoid biosynthesis pathway is relatively well studied, the genetic regulation of glyceollin induction is unclear. Particularly, the contribution, prevalence, and occurrence of gene clusters in plant metabolic diversity are largely unclear. Our mGWAS results suggest there are two gene clusters with functionally related but non-homologous genes, which may involve in glyceollin induction in wild soybean. Thus far, these are the first reported plausible gene clusters involved in glyceollin accumulation induced by biotic stimuli. These gene clusters suggest that glyceollin may be synthesized where the enzyme-encoding genes are adjacent to each other on the same chromosome (Chavali and Rhee, 2018). Physical clustering of genes with similar functions can facilitate co-inheritance of alleles with favorable combinations and their coordinated regulations at chromatin level (Chu et al., 2011, Osbourn, 2010a). Besides, such clusters incline to locate in the sub-telomeric regions (Gierl and Frey, 2001, Qi et al., 2004, Sakamoto et al., 2004), near the ends of chromosomes that are known to harbor mutations. For example, an examination of the complete genome sequence revealed that the maize *DIMBOA* cluster is located close to the end of chromosome 4 (Farman, 2007, Jonczyk et al., 2008). Thus, identifying the positions of the genes can contribute to inferences of possible mechanisms underlying chemical diversity in natural populations.

Tailoring enzymes, such as methyltransferases, glycosyltransferases, *CYPs*,dehydrogenases/reductases, and acyltransferases are responsible for modifying the chemical backbone of specialized metabolites (Osbourn, 2010b). The gene clusters we found are associated with tailoring or regulating glycosyltransferase enzymes. A common defense mechanism of plants involves glycosylation of secondary metabolites by involving these enzymes (Mylona et al., 2008). Therefore, the clustering of the genes encoding glycosyltransferase on chromosome 9 indicates the formation of stress-induced (i.e., SCN stress in our study) protective compounds. For example, the cyclic hydroxamic acid (*DIBOA*) in maize (Frey et al., 1997, Gierl and Frey, 2001), the triterpene avenacin in oat (Field and Osbourn, 2008, Mugford et al., 2009, Qi et al., 2004, Qi et al., 2006), and two gene clusters associated with diterpene (momilactone and phytocassane) synthesis in rice, which may be pre-formed or synthesized after stress induction for plant defense. Disruption of such gene clusters may compromise pest and disease resistance and lead to the accumulation of toxic pathway intermediates (Chu et al., 2011). In the multi-step plant specialized metabolic pathways, rapid adaptation to a particular environmental niche could result in highly diverse and rapidly evolving metabolic gene clusters (Osbourn and Field, 2009). Hence, the level of conservation of the identified gene clusters in this study across different *Glycine soja* genotypes can shed light on evolutionary insight of these clusters (Field and Osbourn, 2008). Synthetic biology and functional genetics can further help investigate the organization and contribution of these clusters in metabolite diversity, as well as decipher the mechanism of adaptive evolution and genome plasticity (Chu et al., 2011, Osbourn, 2010b).

### Plausible transcriptional factors in glyceollin induction

Advancement of genetics, genomics, and bioinformatic approaches facilitate the prediction and identification of a large number of genes, including transcription factors associated with plant-specialized metabolic pathways (Anarat-Cappillino and Sattely, 2014, Moore et al., 2019). However, the transcriptional regulators of specialized metabolism are less well characterized (Shoji and Yuan, 2021). The regulation of highly diverse plant specialized metabolic pathways is dynamic given the ever-changing environment. Such regulation generally occurs at transcription level, and thus, it requires coordinated regulation often mediated by transcription factors (TFs) (Colinas and Goossens, 2018, Shoji, 2019). For instance,*MYB* and basic helix-loop-helix (*bHLH*)TF family genes were reported to regulate anthocyanin and related flavonoid biosynthetic pathways in a wide range of species (Chezem and Clay, 2016). Moreover, significant modifications of these regulatory genes give rise to the vast diversity in plant specialized metabolism (Huang et al., 2018, Springer et al., 2019).

It is possible that transcription factors, such as *MYB* and *WRKY* TFs on chromosome 9, may influence glyceollin induction. This indicates regulation of glyceollin induction with SCN stress may involve a highly complex interplay among multiple genes and pathways. Previous studies reported that gene families of transcription factors, such as *NAC*, *MYB, bHLH*, and *WRKY*,exhibited conservative patterns among *Arabidopsis*, cotton, grapevine, maize, and rice (Ibraheem et al., 2015, Ogawa et al., 2017, Saga et al., 2012, Xu et al., 2004, Yamamura et al., 2015, Zheng et al., 2006). These plant species produce various phytoalexins, such as indole alkaloids, terpenoid aldehydes, stilbenoids, deoxyanthocyanidins, and momilactones/ phytocassanes, respectively. This gives rise to the question of whether these TFs are as diversified as the metabolic pathways, or they maintain conservative patterns among species. The investigation of TFs binding promoter regions can give insights if the pathways are co-opted into stress-inducible regulation by the respective TFs (Jahan et al., 2019). The homology of TFs among different plant species can help metabolic engineering a wide variety of crop plants to produce phytoalexins in greater amounts.

In addition to enzyme-encoding genes, TF genes can also be found as gene clusters. For example, the gene cluster of TF *ERF* (jasmonate (JA)-responsive ethylene response factor) consists of five *ERF* genes in tomato (Cárdenas et al., 2016, Thagun et al., 2016), while eight in potato (Cárdenas et al., 2016), two clusters of ten and five in tobacco (Kajikawa et al., 2017), five in *C. roseus* (Singh et al., 2020), four in *Calotropis gigantea* (Singh et al., 2020), and four in *Glesemium sempervirens* (Singh et al., 2020). Besides, TFs involved in plant specialized metabolism can be found in arrays (Shoji and Yuan, 2021, Zhou et al., 2016). So, it is possible that the TFs we identified are located in the same genomic neighborhood as arrays or biosynthetic gene clusters (BGCs). The co-regulation hypothesis of gene clusters poses that clustering of TFs can help coregulate genes in a pathway. Although co-regulation also exists between un-clustered metabolic pathways, clustering may accelerate the recruitment of genes into a regulon (Smit and Lichman, 2022, Wisecaver et al., 2017).

### Epistasis and plausible selection on glyceollin induction

Metabolic traits have been reported to have low heritability due to environmental effects on their accumulations (Rowe et al., 2008). Recent studies have shown strong epistatic interactions of genes influencing variation of plant specialized metabolites, which may impact fitness in the field (Brachi et al., 2015, Kerwin et al., 2015, Kerwin et al., 2017). For example, numerous epistatic interactions influence the highly complex genetic architecture responsible for *Arabidopsis* metabolism (Kliebenstein, 2001, Kliebenstein et al., 2001). Moreover, a mixture of positive and negative epistatic interactions can help identify significant QTLs located within a biosynthetic pathway (Rowe et al., 2008). Compared to expression regulations, the power of epistasis in metabolomics is that they can better indicate the interconnectedness of metabolites within the metabolic pathway (Arita, 2004, Fell and Wagner, 2000, Jeong et al., 2000). The widespread interactive effects found among our identified significant SNPs affecting targeted metabolic traits may be a consequence of the interconvertibility between daidzein and glyceollin.

Genes containing causal variation for plant defensive compounds may influence field fitness and thus are likely under natural selection (Kroymann, 2011). For example, Benderoth et al. (2006) detected positive selection in glucosinolate diversification in *Arabidopsis thaliana* and its relatives (Benderoth et al., 2006). Prasad et al. (2012) showed positive selection for a mutation on a metabolic pathway gene could enhance resistance to herbivory in natural populations of a rocky mountain cress species (Prasad et al., 2012). We detected strong signals of selection on the SNPs significantly associated with glyceollin phenotypes with EHH and LD analyses (Figs **4**, **2b**,and **S1**). For example, the LD surrounding the significant SNP ss715603454 that is next to the identified gene clusters is more extensive, suggesting strong selection in this region (Figs **2b**, **S1**). Meanwhile, the two alleles of this significant SNP, G and T, showed different EHH values, with T exhibiting much longer haplotype homozygosity. This indicates that this T allele may be under recent positive selection. Interestingly, the T allele is significantly associated with higher induction of glyceollin and has a higher frequency in South Korea (Fig. **2c,d**). The allele-specific EHH pattern and their geographic distribution may be due to heterogeneous selection pressure in nature.

### Perspectives and future directions of our study

Plant specialized metabolites exhibit extreme quantitative and qualitative variation. Therefore, high-throughput metabolite profiling, such as LC-MS analysis coupled with GWAS (as applied here) can help better understand the genetic contributions to metabolic diversity in natural populations. A common assumption is that biological variables or traits should show a normal distribution, and skewed data may indicate measurement error. However, the scenario is different in metabolomics, especially in secondary metabolism. For instance, a ratio of two related compounds, rather than their separate values, may provide a comprehensive understanding of the underlying enzymatic process (Byrne et al., 1996, Chan et al., 2011, Kliebenstein, 2001, Kliebenstein et al., 2001, Kliebenstein, 2007, McMullen et al., 1998, Petersen et al., 2012, Prasad et al., 2012, Yencho et al., 1998). We used a ratio of glyceollin and daidzein concentrations as the phenotypic trait for our association study. The use of a metabolic ratio also may produce: (1) a reduction in the variability of the data collected for the biological replicates and thus increase statistical power and (2) a reduction in overall noise in the dataset by canceling out systemic experimental errors. Most importantly for our purposes, the glyceollin to daidzein metabolite ratio is correlated to the corresponding reaction rate under optimal steady-state assumptions, as this metabolite pair is connected in the phenylpropanoid biosynthetic pathway (Petersen et al., 2012, Suhre et al., 2011).

The natural world has a lot to offer in tackling diseases and global food scarcity. There is a need to develop new medicines and future value-increased food by unlocking the uncharted gene pools of wild plants. Our chosen study system crop wild relative of soybean poses much higher and underexplored genetic diversity than its domesticated descendants. Given that glyceollin is produced in trace amounts, it is an exciting challenge to define the plant metabolic gene clusters and transcriptional regulators in the glyceollin biosynthesis pathway. Besides complex cancer treatment and therapies, the rise of different types of tumors and tumor subtypes urges the need for new drugs. Along with glyceollin’s role in plant defense, it has been well-documented for anti-cancer activities. Our follow-up studies will apply transcriptomics and functional validation of the candidate genes, which can expand our focus to explore associations of genes in clusters to understand their involvement in regulating glyceollin biosynthesis at the systems level. As phytochemical variation can be caused by both structural genes and their expression differences, it will be interesting to explore the role of pathway-specific regulators (i.e., transcription factors) in glyceollin induction (Osbourn, 2010b). Our results suggest that improving our fundamental knowledge of plant specialized metabolic gene clusters and regulators will facilitate metabolic engineering with improved metabolic traits for sustainable agriculture and novel pharmaceuticals.

## Abbreviations list

bp: base pair
BLINK: bayesian-information and linkagedisequilibrium iteratively nested keyway
dpi: days post infection
FDR: false discovery rate
Fig. (Figs): figure (figures)
LD: linkage disequilibrium
LOD: logarithm of the odds
Mbp: megabase pair
mGWAS: metabolite-based genome-wide association study
SNP: single nucleotide polymorphism
μm: micromolar
μg/g: microgram/gram

## Acknowledgements

Research in B-H.S. lab was supported by the National Institute of General Medical Sciences, Award Number: R15GM122029, R15AT011603; North Carolina Biotechnology Center, Award Number: 2019-BIG-6507, 2020-FLG-3806; The North Carolina Soybean Producers Association, Award Number: 18-0252; and the University of North Carolina at Charlotte, Award Number: 111239. Work by F.Y. was (partially) supported by the Schlumberger Foundation Faculty for the Future program.

We thank Thomas Mitchell-Olds for the constructive comments and suggestions. We thank Xu Li and Janice Kofsky for their help with methods and discussions. We also thank Song lab members Melissa Hatley, Judy Janiol, Janice Kofsky and Neha Mittal for their help with planting and sample collection.

## Author contributions

B-H.S. conceived the study and designed the experiment. H.Z., J.W., R.R., C.B. and F.Y. conducted the experiments and collected the data. F.Y., H.Z., L.L., and B.W. performed data analysis. F.Y. drafted the original manuscript with input from. F.Y., H.Z., L.L., B.W., J.W. and B-H.S. reviewed and edited the manuscript. All authors have read and agreed to the published version of the manuscript.

## Notes

### Competing Interest Statement

The authors have declared no competing interest.

### Summary of Updates

The section on "Metabolite extraction and quantification" (Line 150-153) was updated to delete repeated information and provide service location. The section "Acknowledgment" was updated to add Dr. Xu Li.

