## Supplemental_Figures_Tables for "Genetic basis and selection of glyceollin induction in wild soybean" (from another version of this preprint: version 1, 2022-12-19)

### Data availability

### Supporting information

The following Supporting Information is available for this article:

**Fig. S1** A pairwise linkage disequilibrium between target SNP (ss715603454) and candidate gene *Glyma.09G128200* (location of the gene is indicated as a gray hexagon) from our metabolic gene cluster of interest.

**Fig. S2** Significant SNPs ss715585948 **(a)**, ss715615975 **(b)**, ss715620269 **(c)**, and ss715636844 **(d)** on chromosomes 3, 13, 15, and 20, respectively, show narrow LD blocks.

**Table S1** Wild soybean ecotypes used in this study. (MS Excel file)

**Table S2** SNP effect and heritability estimate for the significant SNPs related to plant specialized metabolic pathway.

**Table S3** Gene of interest with enzyme class and associated metabolic domain for chromosome 9.

**Table S4** Annotation of candidate genes of gene clusters 1 and 2.

**Table S5** Annotation of candidate genes other than genes within cluster 1 and 2.

**Fig. S1** A pairwise linkage disequilibrium between target SNP (ss715603454) and candidate gene *Glyma.09G128200* (location of the gene is indicated as a gray hexagon) from our metabolic gene cluster of interest.

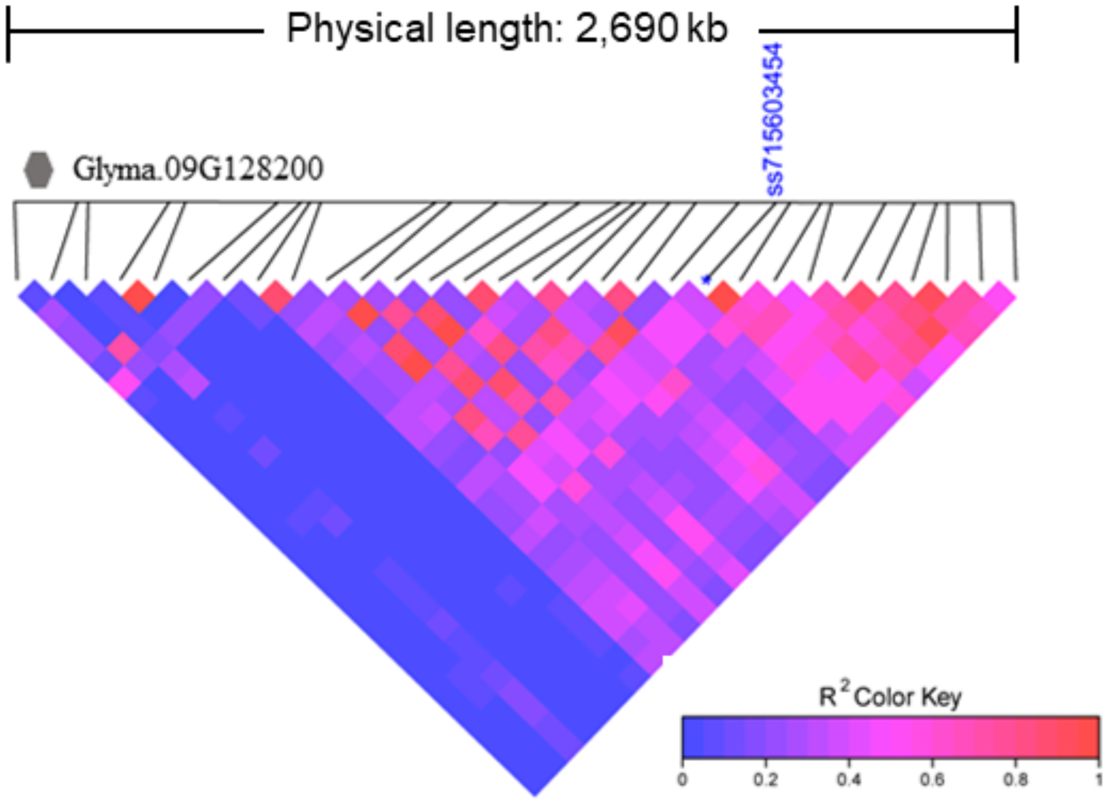

**Fig. S2** Significant SNPs ss715585948 **(a)**, ss715615975 **(b)**, ss715620269 **(c)**, and ss715636844 **(d)** on chromosomes 3, 13, 15, and 20, respectively, show narrow LD blocks.

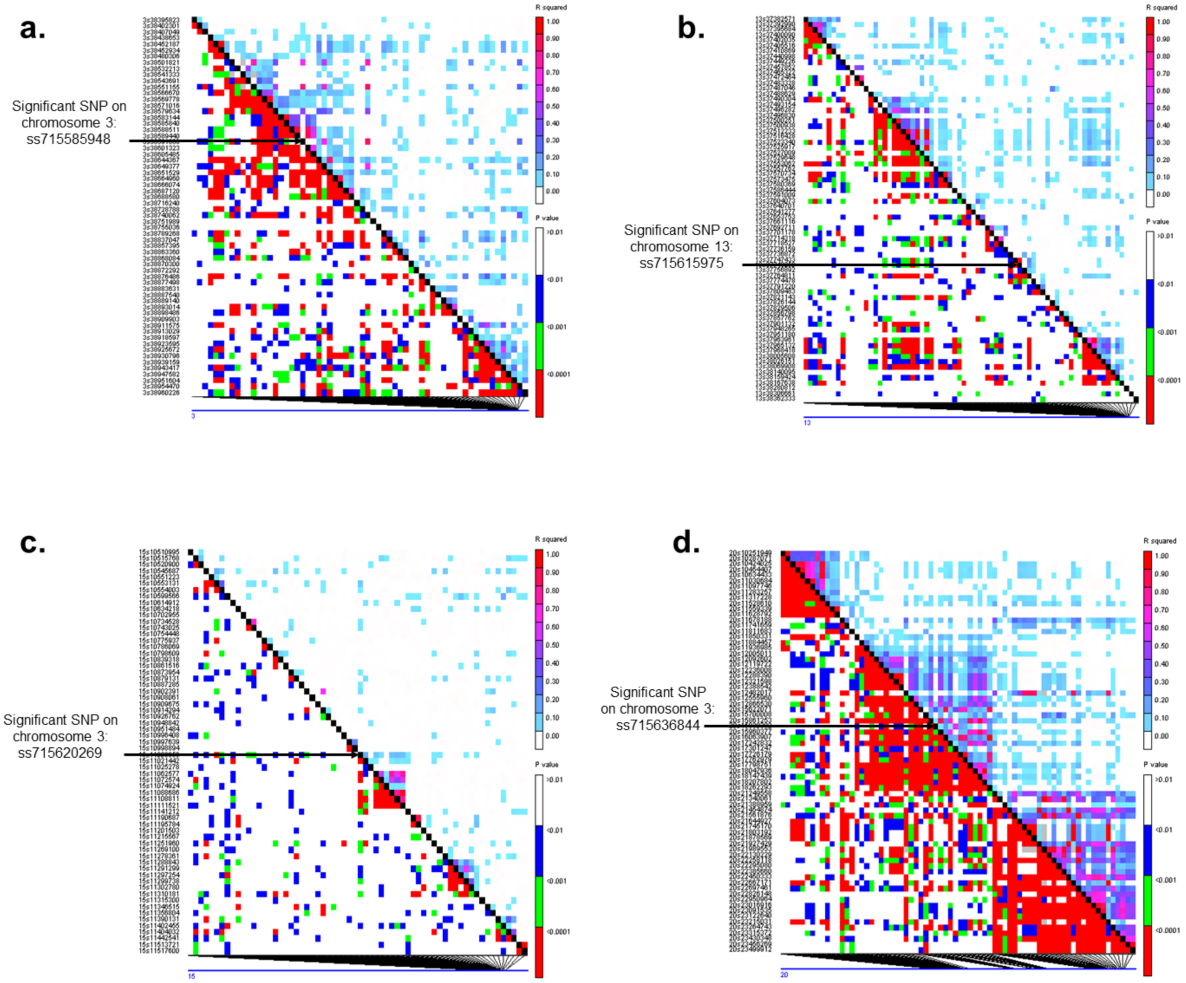

**Table S1** Wild soybean ecotypes used in this study. (MS Excel file)

**Table S2** SNP effect and heritability estimate for the significant SNPs related to plant specialized metabolic pathway.

| **Significant SNP** | **Chromosome** | **Position** | **LOD score** | **SNP effect** | **% R-squared value** | **$h^2$ (%)** |
|---|---|---|---|---|---|---|
| ss715585948 | Gm03 | 38591888 | 3.80 | -0.31 | 10.58 | 35 |
| ss715603454 | Gm09 | 30262482 | 3.80 | 0.06 | 4.22 | |
| ss715603455 | Gm09 | 30191235 | 3.80 | 0.05 | 3.32 | |
| ss715603462 | Gm09 | 30393285 | 3.80 | -0.06 | 3.96 | |
| ss715603471 | Gm09 | 30725658 | 3.80 | -0.06 | 3.86 | |
| ss715615975 | Gm13 | 37748250 | 3.81 | 0.22 | 12.59 | |
| ss715620269 | Gm15 | 11003656 | 3.81 | -0.20 | 11.42 | |
| ss715636844 | Gm20 | 15930392 | 3.79 | -0.13 | 4.16 | |

Locations of the significant SNPs shown in base pairs, bp; standardized SNP effects, and the percentage (%) of the total phenotypic variation (glyceollin induction) explained by significant SNPs (% R- squared values) on different chromosomes. LOD scores represent the chromosome-wide significant level from 3.79 to 3.82. Heritability ($h^2$) of glyceollin induction was calculated with all SNPs.

**Table S3** Gene of interest with enzyme class and associated metabolic domain for chromosome 9.

| **Gene Cluster** | **Gene Name** | **Enzyme Class** | **Signature or Tailoring?** | **Metabolic Domain** |
|---|---|---|---|---|
| **Cluster 1** | *Glyma.09G127200*<br>*Glyma.09G127300* | glycosyltransferase | tailoring | Phenylpropanoid Derivatives; |

| | | | |
|---|---|---|---|
| | | | Specialized Metabolism |
| **Cluster 2** | *Glyma.09G127700*<br>*Glyma.09G128200*<br>*Glyma.09G128300*<br>*Glyma.09G128400* | glycosyltransferase tailoring | Phenylpropanoid Derivatives; Specialized Metabolism |

**Table S4** Annotation of candidate genes of gene clusters 1 and 2.

| **Genes** | **Annotation** |
|---|---|
| *Glyma.09G127200* (cluster 1) | *UGT88A1* (AT3G16520.3); isoflavone 7-O-glucosyltransferase (2.4.1.170); metabolic process (GO:0008152); hexosyltransferase activity (GO:0016758), glucosyl/glucuronosyl transferases (PTHR11926); UDP-glucoronosyl and UDP-glucosyl transferase (PF00201) |
| *Glyma.09G127300* (cluster 1) | *UGT88A1* (AT3G16520.3); isoflavone 7-O-glucosyltransferase (2.4.1.170); metabolic process (GO:0008152); hexosyltransferase activity (GO:0016758); glucosyl/glucuronosyl transferases (PTHR11926); UDP-glucoronosyl and UDP-glucosyl transferase (PF00201); glycosyltransferases (K08237) |
| *Glyma.09G127700* (cluster 2) | *UGT88A1* (AT3G16520.3); isoflavone 7-O-glucosyltransferase (2.4.1.170); metabolic process (GO:0008152); hexosyltransferase activity (GO:0016758), glucosyl/glucuronosyl transferases (PTHR11926); UDP-glucoronosyl and UDP-glucosyl transferase (PF00201) |
| *Glyma.09G128300* (cluster 2) | *UGT88A1* (AT3G16520.3); isoflavone 7-O-glucosyltransferase (2.4.1.170); metabolic process (GO:0008152); hexosyltransferase activity (GO:0016758), glucosyl/glucuronosyl transferases (PTHR11926); UDP-glucoronosyl and UDP-glucosyl transferase (PF00201) |

| | |
|---|---|
| *Glyma.09G128200* (cluster 2) | UDP-glucosyl transferase 88A1 (AT3G16520.3); metabolic process (GO:0008152); intracellular membrane-bounded organelle (GO:0043231); transferase activity; transferring hexosyl groups (GO:0016758); quercetin 3-O-glucosyltransferase activity (GO:0080043); quercetin 7-O-glucosyltransferase activity (PWY-2345); biochanin A conjugates interconversion (PWY-2861); isoflavone 7-O-glucosyltransferase (GN7V-57685) |
| *Glyma.09G128400* (cluster 2) | *UGT88A1* (AT3G16520.3), metabolic process (GO:0008152), hexosyltransferase activity (GO:0016758), glucosyl/glucuronosyl transferases (PTHR11926), and UDP-glucoronosyl and UDP-glucosyl transferase (PF00201) |

**Table S5** Annotation of candidate genes other than genes within cluster 1 and 2.

| **Genes** | **Annotation** |
|---|---|
| *Glyma.20G052000, Glyma.20G052400* | UDP-Glycosyltransferase superfamily protein |
| *Glyma.20G057500* | UDP-glucosyl transferase 85A2 |
| *Glyma.20G058000* | hydroxy methylglutaryl CoA reductase 1 (Mevalonate pathway I; isoprenoid biosynthetic process; sterol biosynthetic process; coumarin biosynthetic process; oxidoreductase activity) |
| *Glyma.20G053900* | *cytochrome P450*, family 71, subfamily B, polypeptide 34 (Oxidoreductase activity) |
| *Glyma.20G065000, Glyma.20G065100* | *cytochrome p450* 79a2 |
| *Glyma.13G272500* | *bZIP* transcription factor |
| *Glyma.09G125400, Glyma.13G272700, Glyma.13G273500, Glyma.13G277600,* | *RING/U-box* superfamily protein, *RING/FYVE/PHD* zinc finger superfamily protein |

| | |
|---|---|
| *Glyma.15G134700,*<br>*Glyma.20G064600* | |
| *Glyma.03G176600,*<br>*Glyma.09G129100,*<br>*Glyma.09G127100*<br>*Glyma.15G139000,*<br>*Glyma.15G135600* | *WRKY* family transcription factor family protein |
| *Glyma.09G113000,*<br>*Glyma.09G113100,*<br>*Glyma.15G134100* | *myb* domain |
| *Glyma.03G173100,*<br>*Glyma.03G173200,*<br>*Glyma.03G173300,*<br>*Glyma.09G128700,*<br>*Glyma.09G115100,*<br>*Glyma.09G107400*<br>*Glyma.13G274600,*<br>*Glyma.20G059300* | zinc fingers superfamily protein |
| *Glyma.13G274300,*<br>*Glyma.13G279900,*<br>*Glyma.13G280000* | *NAC* transcription factors |
| *Glyma.09G115900,*<br>*Glyma.09G117400,*<br>*Glyma.09G123200,*<br>*Glyma.13G277100,*<br>*Glyma.20G053900,*<br>*Glyma.20G065000,*<br>*Glyma.20G065100* | *cytochrome P450* enzyme family |
| *Glyma.20G065000,*<br>*Glyma.20G065100* | *cytochrome p450* 79a2 (oxidoreductase activity; Linamarin biosynthesis**)** |

| | |
|---|---|
| *Glyma.09G108800* | Zinc finger, *RING-type*; Transcription factor jumonji/aspartyl beta-hydroxylase |
| *Glyma.09G109500, Glyma.09G122500* | Terpenoid cyclases family protein, terpene synthase 03 |
| *Glyma.03G176300* | Glutathione S-transferase family protein (glucosinolate biosynthetic process) |
| *Glyma.09G126500 Glyma.09G126900 Glyma.15G139200* | phenylpropanoid metabolic process |
